# Orbitofrontal cortex promotes trial-by-trial learning of risky, but not spatial, biases

**DOI:** 10.1101/685107

**Authors:** Christine M. Constantinople, Alex T. Piet, Peter Bibawi, Athena Akrami, Charles D. Kopec, Carlos D. Brody

**Affiliations:** Princeton Neuroscience Institute, Princeton University, Princeton, NJ, 08544, USA; Department of Molecular Biology, Princeton University, Princeton, NJ, 08544, USA; Howard Hughes Medical Institute, Princeton University, Princeton, NJ, 08544, USA; Center for Neural Science, New York University, NY 10003, USA; Allen Institute for Brain Science, Westlake Avenue N, Seattle, WA, 98109, USA; Sainsbury Wellcome Centre, 25 Howland Street London W1T 4JG

## Abstract

Individual choices are not made in isolation but are embedded in a series of past experiences, decisions, and outcomes. The effects of past experiences on choices, often called sequential biases, are ubiquitous in perceptual and value-based decision-making, but their neural substrates are unclear. We trained rats to choose between explicitly cued guaranteed and probabilistic rewards in a task in which outcomes on each trial were independent. Behavioral variability often reflected the effects of previous trials, including increased willingness to take risks following risky wins, and spatial “win-stay/lose-shift” biases. Electrophysiological recordings from lateral orbitofrontal cortex (lOFC) revealed strong encoding of reward history and receipt, and optogenetic inhibition of lOFC during choice reporting eliminated rats’ increased preference for risk on subsequent trials following risky wins, but spared other sequential effects. Our data show that different sequential biases are neurally dissociable, and that the lOFC’s role in adaptive behavior promotes learning of more abstract, task-specific biases (here, biases for the risky option), but not spatial ones. These data are consistent with proposals that the OFC represents the animal’s location in an abstract representation of task states, and highlight the lOFC’s role in learning.

## Introduction

Sequential biases permeate human decision-making, and while using past experiences to guide decision-making can be useful in dynamic environments, it can cause us to make suboptimal decisions if the past is not informative (but see Yu and Cohen, 2008). A variety of biases have been described in two-alternative forced choice tasks in humans and animal models, including repetition of successful choices (“win-stay”), switching after unsuccessful choices (“lose-switch”), and biases due to previous sensory experiences (Abrahamyan et al., 2016; Akrami et al., 2018; Busse et al., 2011; Hollingworth, 1910; Kagel et al., 1995; Scott et al., 2015; Visscher et al., 2009). Moreover, depending on task design, these biases can be expressed in different ways: if choice options are fixed in space, as is often the case in studies of rodent behavior, sequential biases will appear spatial and action-dependent. Alternatively, if the choice options are not fixed in space, sequential biases may be expressed in non-spatial, task-dependent coordinates. It is unclear whether different sequential biases, or biases expressed in different coordinate reference frames, rely on shared or distinct neural circuits and mechanisms.

We trained rats to choose between explicitly cued guaranteed or probabilistic (i.e., risky) rewards (Constantinople et al., 2019). The location of the guaranteed and risky rewards were randomly assigned to different locations on each trial, disambiguating biases expressed in spatial coordinates and those expressed in more abstract, task-specific coordinates (biases for risky or safe options). We found that the lateral orbitofrontal cortex (lOFC) was required for rats’ expression of task-specific, but not spatial, sequential biases. We interpret these data as consistent with proposals that OFC represents the animal’s location in an abstract representation of task states (Wilson et al., 2014).

OFC in mice, rats, monkeys, and humans has been shown to play a critical role in adapting behavior to dynamic task contingencies, especially when those contingencies are partially observable (Bradfield et al., 2015; Rushworth et al., 2011; Stalnaker et al., 2014; Wallis, 2007; Wilson et al., 2014). Lesion experiments have implicated the OFC in tracking rewards and value, for example in extinction, devaluation, and reversal learning paradigms, in which reward contingencies were explicitly manipulated by the experimenter (Gallagher et al., 1999; Izquierdo et al., 2004; Pickens et al., 2003). A related body of work implicates the OFC in evaluative processes, including comparing current choices to previous outcomes (Kennerley et al., 2011), and regret (Steiner and Redish, 2014, 2012).

OFC also represents the subjective value of goods in economic decision-making tasks (Padoa-Schioppa and Conen, 2017). In this context, OFC neurons adapt their firing rates to the range of rewards the animal experiences (Conen and Padoa-Schioppa, 2019; Kobayashi et al., 2010; Padoa-Schioppa, 2009; Saez et al., 2017; Zimmermann et al., 2018), thereby dynamically encoding subjective value to accurately reflect changes in task contingencies, here, reward statistics.

Most of these studies demonstrating that OFC promotes behavioral flexibility have used tasks in which trial-by-trial learning improves behavioral performance (e.g., reversal learning). We hypothesized that OFC’s role in behavioral flexibility might also drive deleterious sequential biases. We developed a task in which sequential biases were maladaptive: trials were independent and reward contingencies were stable over months of training (Constantinople et al., 2019). Rats exhibited several dissociable biases reflecting trial and reward history, and we identified lOFC as critical for one bias in particular: an increased willingness to take risk following risky wins.

## Results

### Rats choosing between guaranteed and probabilistic rewards show a risky “win-stay” bias

We developed a task in which rats chose between explicitly cued guaranteed and probabilistic rewards (Constantinople et al., 2019). Animals initiated a trial by nose-poking in the center of three ports. Auditory clicks were presented from left and right speakers, and the click rate (6-48 Hz) conveyed the volume of water reward baited at each of the two side ports. Simultaneously, light flashes were presented from side ports, and the number of flashes (0-10) conveyed the probability of receiving water reward at each port (Figure 1A). One port offered a guaranteed or “safe” reward (p=1), and the other offered a probabilistic or “risky” reward (p≤1). Rewards were delivered 100ms after rats entered the side port. The location of the safe and risky ports varied randomly on each trial. Four possible water volumes were offered (6, 12, 24, 48μL), and risky reward probabilities ranged from 0 to 1, in increments of 0.1 (Figure 1B,C). The trials were self-paced, and following a choice, rats were free to initiate the next trial within 100-200ms, although they typically took longer (Figure 1 - figure supplement 1). However, if animals terminated the trial prematurely by breaking center fixation, they were penalized with a time-out penalty (1.5-8 seconds, adjusted across rats as needed).

**Figure 1.**
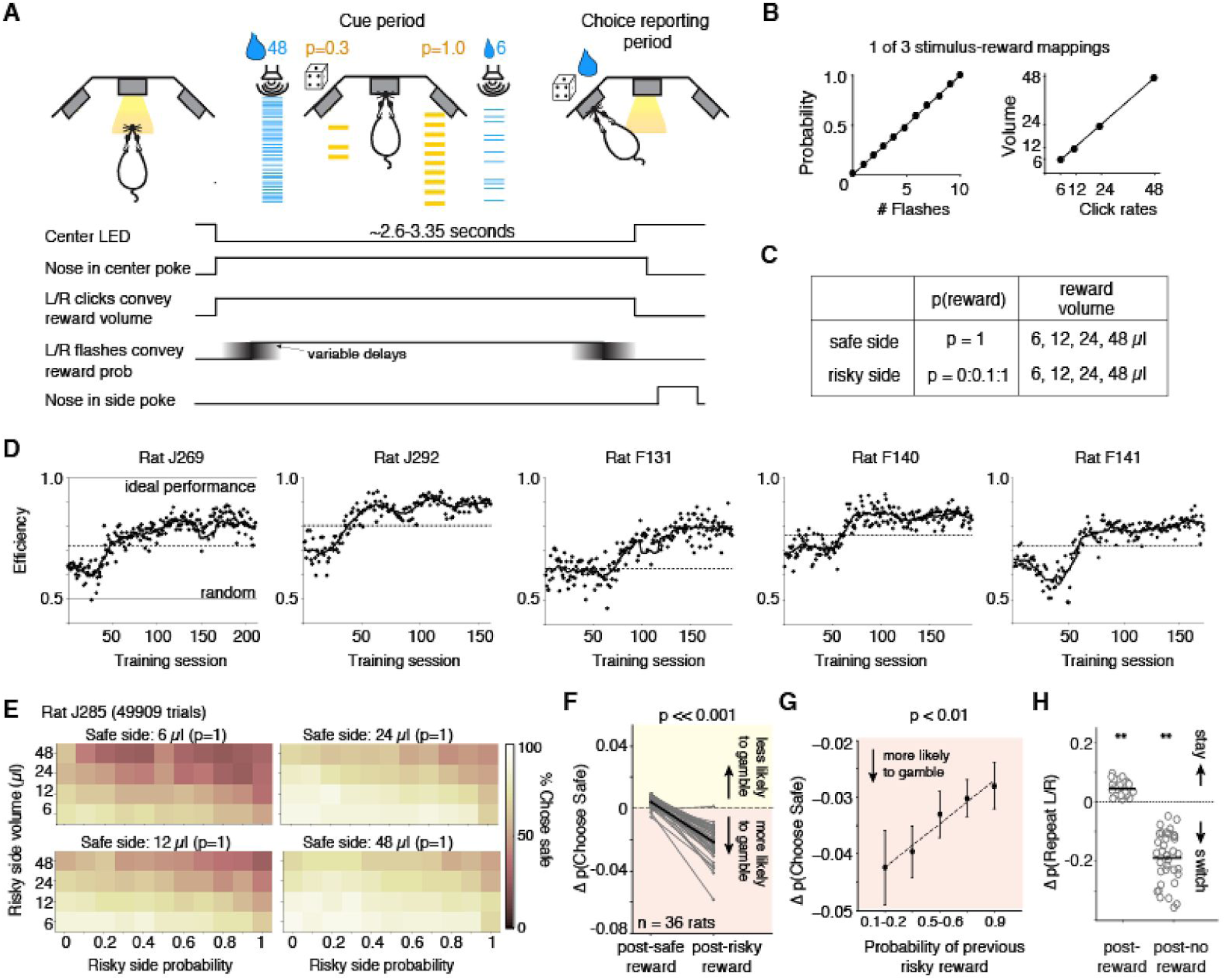
Behavioral task: Rats performing the task exhibit stable performance over months, but also trial-by-trial learning dynamics. (A) Example trial: rat initiates a trial by nose-poking and fixating in center. On each side, light flashes and click rates convey reward probability and water volume, respectively. One side (here, the right port) offers guaranteed reward (“safe”); safe and risky sides vary randomly over trials. (B) Relationship between flashes and probability, and click rates and reward volumes (6, 12, 24, or 48μL) in one version of the task. Risky side could have reward probability between 0-1 (increments of 0.1). (C) Offered reward volumes and probabilities. (D) Behavioral performance in units of “efficiency” for five representative rats in the final training stage (Methods). We compared the average expected value (reward x probability) per trial the rat received compared to an agent choosing randomly, or one that always chose the option with the greater expected value (“ideal performance”). The dashed line is criterion performance for each rat (see “Methods”). (E) Percent of trials one rat chose the safe option for each of the four safe volumes. Axes show probability and volume of risky alternatives. (F) Difference in probability of choosing the safe option following guaranteed rewards and risky rewards (relative to the mean probability of choosing safe) for all rats (black is mean). Rats were more likely to gamble following risky rewards (p=8.35e-16, paired t-test). (G) The magnitude of the risky win-stay bias exhibits graded dependence on the reward probability of the gamble (mean across rats). p=0.0035 of slope parameter of least-squares regression line (dashed line). The riskier the gamble that won, the more likely that rats will choose to gamble again. See also Figure 2 - figure supplement 1. (H) Change in the probability of repeating left or right choices following rewarded or unrewarded trials. Asterisks indicate that rats’ “win-stay” biases were significantly different from zero (p=2.06e-13, paired t-test), as were their “lose-switch” biases (p=2.65e-15).

The “cue period” is the period when the rat is center poking, and flashes and clicks are presented. The “choice reporting period” begins when the rat exits the center poke and is free to report his choice by poking in one of the side ports.

To determine when rats were sufficiently trained to understand the meaning of the cues in the task, we evaluated the “efficiency” of their choices, by comparing their mean expected value (probability x reward) per trial relative to an agent that always chose the option with greater expected value (“ideal performance”) and one that chose randomly (“random”; Figure 1D, see also (Rustichini et al., 2017)). While variable, most rats learned the meaning of the cues within 1-2 months of training in the final training stage (Figure 1D). Well-trained rats (n=36 animals) performed, on average, 368 trials per days (+/- 28 trials, s.e.m.; Figure 1 - figure supplement 1B). They tended to choose the option with the higher expected value; on trials when both sides offered certain rewards, rats chose the larger reward, and when one side offered no reward (p=0), they chose the alternative (Figure 1D,E, Figure 1 - figure supplement 1C; Constantinople et al., 2019).

Rats also exhibited sequential biases observed in primates: if they chose the risky option and were rewarded on the previous trial, they were more likely to gamble and choose the risky side again (Figure 1F; Blanchard et al., 2014; Hayden et al., 2008; Neiman and Loewenstein, 2011a). The change in probability of choosing the risky side following a risky win was significantly different from zero (p=1.29e-13, paired t-test across rats). This bias was not due to rats “un-learning” the meaning of the flashes: there was no change in performance on trials where both sides offered the same reward volume, in which case the better option is the guaranteed reward, indicated by the flashes (p=0.397, paired t-test across rats). The magnitude of the risky win-stay bias depended on the risky reward probability (Figure 1G). We emphasize that in this task, the belief that a risky “win” increases the probability of future wins is a fallacy: outcomes are independent on each trial, by design. This result did not reflect biased estimates of conditional probabilities, often observed in finite sequential data (Figure 1 - figure supplement 1D,E). There was no change in rats’ willingness to choose the gamble following a risky loss, indicating that rats were not simply going through phases of preference for risky or safe options (p=0.79 paired t-test; Figure 1 - figure supplement 1F). Similarly, there was no systematic change in rats’ risk preferences over the course of the training session: we examined trials where the guaranteed and risky reward had the same expected value. Rats’ choices on these trials indicate their risk attitudes, with risk averse rats preferring the guaranteed reward and risk seeking rats preferring the gamble (Constantinople et al., 2019). There was no significant change in the probability of choosing the safe option on these trials, comparing the first and second half of trials in each session (p=0.15, paired t-test), again indicating that the risky win-stay bias was not due to slow fluctuations in rats’ risk preferences. Finally, because the risky and safe ports switch on each trial, this risky win-stay bias was independent of a spatial win-stay/lose-switch bias for the left or right ports (Figure 1H), or repetitive behavioral patterns such as perseveration (Miller et al., 2017).

### OFC represents reward history during the cue period

We performed tetrode recordings in lOFC while rats performed the task. Many lOFC neurons exhibited transient responses at trial initiation, the magnitude of which reflected whether the previous trial was rewarded (Figure 2A). To quantify this, we measured the mean discriminability (*d’*) of firing rates at each time point comparing trials following rewarded and unrewarded choices (Figure 2B), analyzing cells with significantly different spike counts on these trials (n=512 of 1459 units; p<0.05, unpaired t-test). lOFC neurons reflected reward history most strongly at trial initiation, when the animal has no information yet about the prospects on the current trial (Figure 2B; Figure 2 - figure supplement 1; (Nogueira et al., 2017). A significantly greater fraction of neurons exhibited higher firing rates following unrewarded trials, compared to rewarded trials, consistent with adaptation effects (Figure 2C; Conen and Padoa-Schioppa, 2019; Kennerley et al., 2011; Padoa-Schioppa, 2009; Saez et al., 2017; Zimmermann et al., 2018).

**Figure 2.**
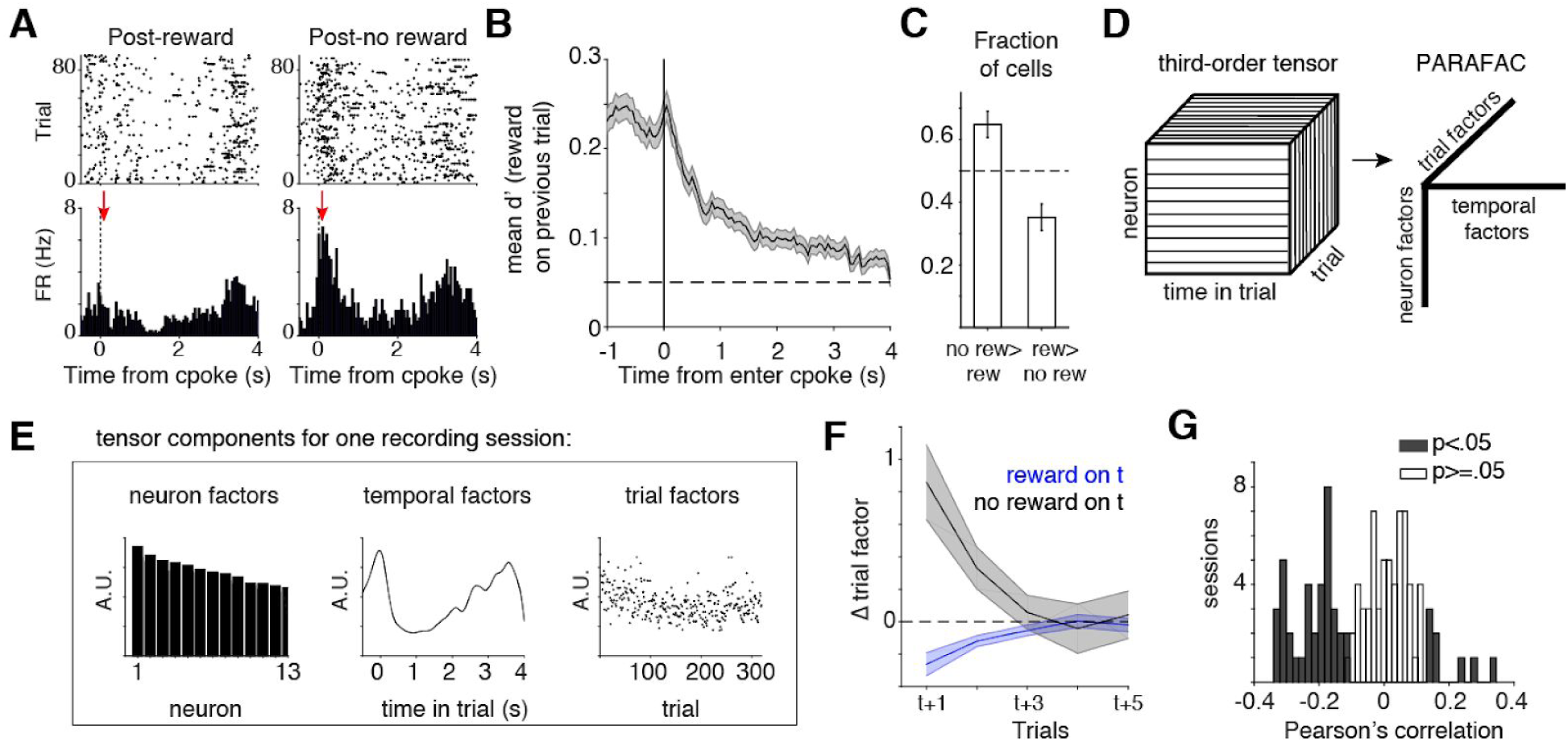
lOFC encodes reward history during the cue period. (A) lOFC neuron with activity aligned to trial initiation. This neuron’s firing rate reflected whether the previous trial was rewarded. (B) Mean encoding of reward history (discriminability or d’) across lOFC neurons that exhibited significantly different spike counts based on reward history. Mean +/- s.e.m. See also Figure 2 - figure supplement 1. (C) Fraction of neurons with significantly different spike counts based on reward history, with more spikes following unrewarded (no rew > rew) or rewarded (rew > no rew) trials. (D) Schematic of analysis (TCA/PARAFAC) used to discover low dimensional descriptions of trial-by-trial population dynamics. See also Figure S3. (E) Result of TCA/PARAFAC from one recording session. Y-axis is in arbitrary units (A.U.; see Methods). (F) Mean (+/- s.d.) shuffle-corrected reward (blue) and no-reward (black) triggered averages of trial factors across all sessions (see Methods). (G) Correlation between trial factors and reward history for each session. Grey bars indicate significance.

Our data show that lOFC neurons encode information about reward history (Figure 2A,B), which might be expected from neurons mediating trial history biases. Given that behavior likely requires the activity of populations of neurons, we next sought an unsupervised description of simultaneously recorded neurons. We employed an extension of principal components analysis, tensor components analysis (Williams et al., 2018); TCA, also known as CANDECOMP/PARAFAC tensor decomposition; Figure 3D). This method extracts features of three aspects of neural data: (1) neuron factors, which weight each neuron’s activity; (2) temporal factors, which capture time-varying dynamics within a trial; and (3) trial factors, which capture dynamics across trials. TCA/PARAFAC provides a low dimensional description of neural dynamics both *within* and *across* trials, allowing for simple descriptions of complex, multi-neuron responses across multiple timescales. TCA provides a key advantage over more common dimensionality reduction techniques like PCA or Factor Analysis which require averaging over trials; TCA allows us to independently quantify trial to trial fluctuations in neural activity. TCA decomposes a 3rd order data tensor *X*_*n,t,k*_ (with n neurons over k trials of length t) by a sum of rank 1 factors 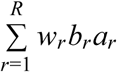. Here, for each rank r, w is a vector of neuron factors, b is a vector of temporal (time within trial) factors, and a is a vector of across trial factors. Figure 2E shows the neuron, temporal, and trial factors for one recording session (see Methods, Figure 2 - figure supplement 2).

**Figure 3.**
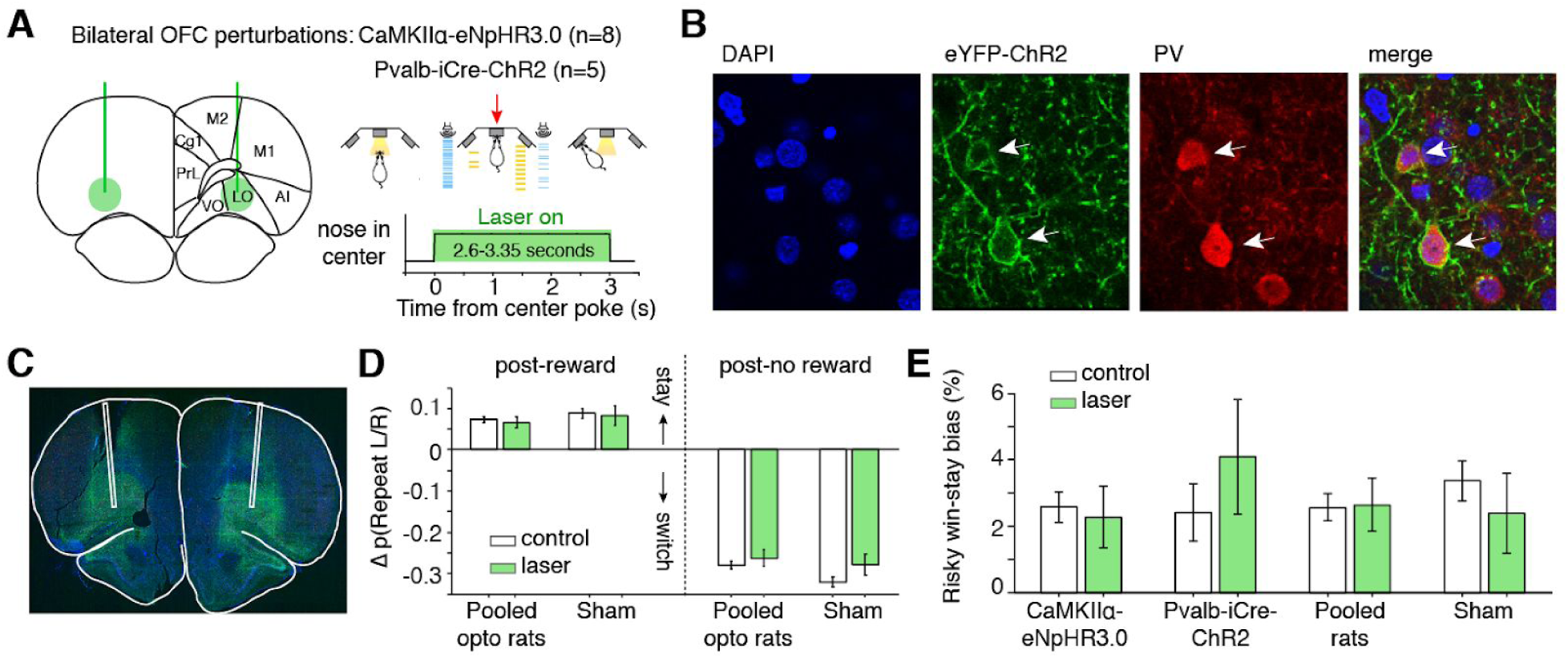
Optogenetic perturbation of lOFC during the cue period does not affect spatial or risky trial history biases. (A) Schematic of bilateral optogenetic perturbations. For CaMKIIα-eNpHR3.0 rats (n=8), we used continuous illumination of a green laser for photoinhibition. For Pvalb-iCre-ChR2 rats (n=5), a blue laser was pulsed at 20Hz. See also Figure S4. While the schematic shows a 3-second trial, trial durations were variable (2.6-3.35 seconds); photoinhibition persisted for the duration of the cue period. (B) Histological section from Pvalb-iCre-ChR2 rats also stained for DAPI and parvalbumin (PV) immunoreactivity. (C) Virus injection in a wild type rat expressing CaMKIIα-eNpHR3.0. Location of fibers were estimated by damage at brain surface and fiber tracks. (D) Magnitude of spatial win-stay and lose-switch biases (difference in probability of repeating a left or right choice) on control and laser trials. Error bars are normal approximation of 95% confidence intervals (Methods). (E) Magnitude of risky win-stay bias (difference in probability of choosing the safe option following safe or risky rewards) on control and laser trials.

Trial factors were significantly modulated by reward history two trials in the past, on average (Figure 2F). This modulation required the neurons whose firing rates reflected reward history (Figure 2 - figure supplement 2E-F). On 43% of recording sessions (45/105 sessions), there was a significant correlation between rats’ reward history and the trial factors, and these correlations were typically negative (Figure 2G). The trial factor can be thought of as a gain factor applied to the population response (Figure 2 - figure supplement 2D); a negative correlation suggests that when rats received reward, neural activity in lOFC generally decreased on the subsequent trial, consistent with several reports of reward adaptation in OFC (Conen and Padoa-Schioppa, 2019; Kennerley et al., 2011; Padoa-Schioppa, 2009; Saez et al., 2017; Zimmermann et al., 2018).

Given the strong encoding of aggregated reward history, we hypothesized that disrupting lOFC dynamics during the cue period would disrupt trial-by-trial learning. Optogenetic inhibition during the cue period, however, did not affect spatial win-stay or lose-switch biases (Figure 3A-D; Figure 3 - figure supplement 1). Photoinhibition also did not affect the risky win-stay bias, in wild-type CaMKIIα-eNpHR3.0 or Pvalb-iCre-ChR2 rats (Figure 3E; p=0.005, one-way ANOVA comparing safe choices following safe or risky rewards, pooling data across all optogenetic rats).

### Disrupting lOFC during the choice report eliminated the risky win-stay bias

In contrast to during the cue period, activity at the time of the choice report, when rats exited the center poke, often reflected whether rats chose the safe or risky prospect (Figure 4A,B). A subset of lOFC neurons exhibited significantly different spike counts on rewarded trials when rats chose the safe compared to the risky option (Figure 4B-E; n=128 units, unpaired t-test). For these neurons, discriminability peaked shortly after rats left the center poke (Figure 4C,D). We also observed prominent side-selectivity (n=628 units) and strong encoding of reward receipt (n=459) during the choice reporting period (Figure 4F-H). Of the neurons whose activity reflected safe/risky choice, there was no significant side bias, and the majority of those units were not selective for left/right choice (Figure 4 - figure supplement 1).

**Figure 4.**
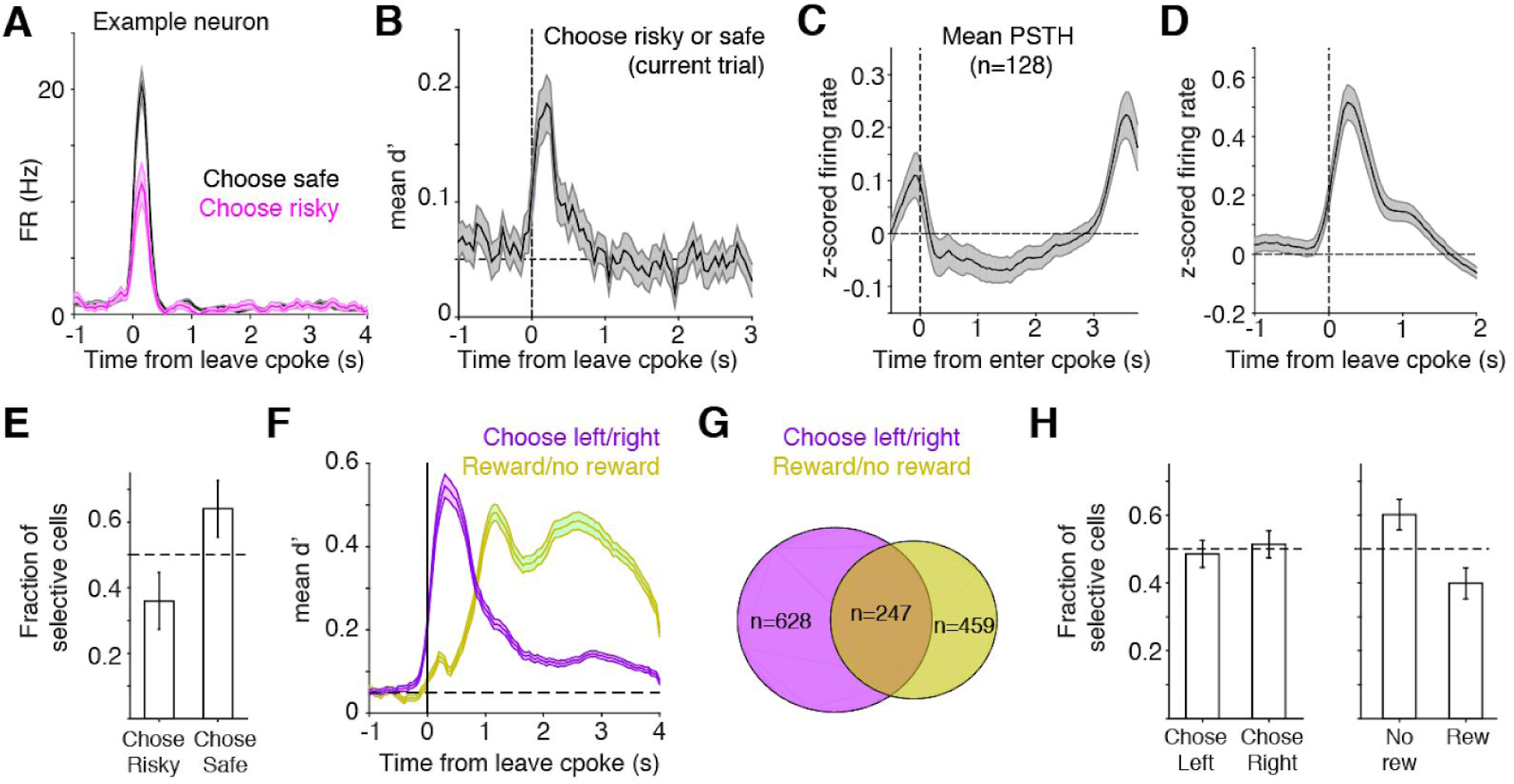
At time of choice report, lOFC neurons represent risk, reward, and left/right choice. (A) Example lOFC neuron with activity aligned to when the rat left the center poke to report his choice. This neuron’s firing rate reflected whether the rat chose the risky (magenta) or safe (black) option on the current trial, analyzing rewarded trials only. (B) Mean d’ across lOFC neurons with significantly different spikes counts on trials with risky or safe choices. See also Figure 4 - figure supplement 1. (C,D) Mean z-scored firing rate of neurons in panel B aligned to entering the center poke (C), or leaving it to report choice (D). (E) Fraction of neurons in panels B-D that preferred trials when rats made risky or safe choices. Higher firing rates on trials in which rats chose the safe reward could reflect encoding of decision confidence or reward expectation (Lak et al., 2014). (F) Mean d’ reflecting whether rats chose the left/right ports, or whether rats received reward, averaged across neurons with significantly different spike counts on those trials. See also Figure 4 - figure supplement 1. (G) Venn diagram of overlap between neurons whose activity differentiated between left/right choices and rewarded/unrewarded trials. (H) Fraction of neurons in panels F,G preferring left/right choices or rewarded/unrewarded trials.

Given the prominent encoding of rats’ left/right choices during this period (Figure 4F,G), we next tested whether photoinhibition affected spatial win-stay/lose-switch biases for the left/right ports. We optogenetically perturbed lOFC during the choice reporting period (triggered when rats exited the center port) for 4 seconds, and then analyzed performance on subsequent trials. Inhibition during the choice reporting period was interleaved in the same sessions as inhibition during the cue period, in the animals shown in Figure 3. While there was a significant reduction in spatial biases, sham rats also exhibited reduced win-stay/lose-switch biases (Figure 5B). Photoinhibition during the choice reporting period imposed a minimum inter-trial interval (ITI) of 4 seconds (for laser illumination), which was longer than the average ITI (Figure 1 - figure supplement 1A). Comparing the photoinhibition trials to control trials with a minimum ITI of 4 seconds eliminated the reduction in spatial win-stay/lose-switch biases in CaMKIIα-eNpHR3.0, Pvalb-iCre-ChR2, and sham rats (data not shown).

**Figure 5.**
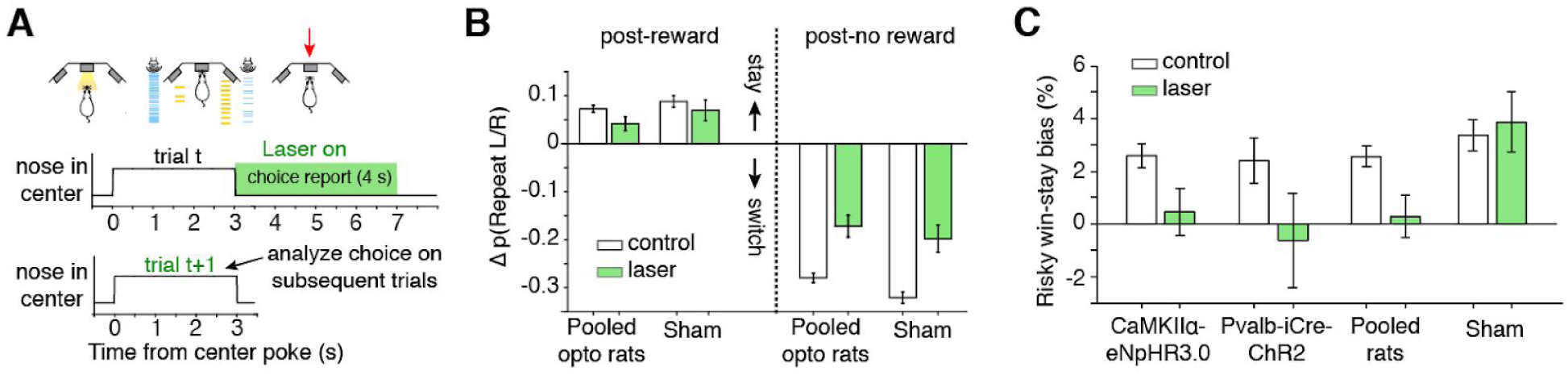
Photoinhibition of lOFC at the time of choice report selectively eliminates the risky win-stay bias. (A) For choice reporting period perturbations, the laser was triggered when rats left the center poke, and persisted for 4 seconds into the inter-trial-interval. See also Figure 5 - figure supplement 1. (B) Spatial win-stay/lose-switch biases following photoinhibition during the choice reporting period; sham rats also exhibited a significant reduction in lose-switch biases, and trended towards a reduction in win-stay biases. (C) Magnitude of the risky win-stay bias following choice reporting period inactivations. Control data are replotted from Figure 4D. Error bars are 95% confidence intervals.

In contrast, optogenetic inhibition of lOFC during the choice reporting period eliminated the risky win-stay bias on the subsequent trial in rats expressing light-sensitive opsins (Figure 5A,C; p=0.667 CaMKIIα-eNpHR3.0; p=0.778 Pvalb-iCre-ChR2; one-way ANOVA comparing safe choices following safe or risky rewards, pooling data across rats). Laser illumination did not disrupt the risky win-stay bias in sham rats not expressing light-activated opsins (Figure 5C; p=0.009, one-way ANOVA comparing safe choices following safe or risky rewards, n=3 rats). For the majority of rats (7/8 CaMKIIα-eNpHR3.0 rats and 4/5 Pvalb-iCre-ChR2 rats), there was no significant effect on choice latencies compared to control trials (Wilcoxon rank-sum test, Bonferroni correction for multiple comparisons). Moreover the behavioral effect occurred on trials *subsequent* to the optogenetic perturbations (Figure 5A), making it unlikely that they were due to off-target illumination of motor cortex, including overlying M2.

We wanted to determine if photoinhibition of lOFC affected other potential biases, so we used logistic regression with parameters for choice hysteresis for safe/risky choices, the risky win-stay bias described above, left/right choice hysteresis, and systematic left/right side biases (Kagel et al., 1995; Padoa-Schioppa, 2013). Photoinhibition during the choice report exclusively reduced the risky win-stay bias parameter, but not others (Figure 5 - figure supplement 1; p=0.0063, one-way ANOVA with Bonferroni correction; Methods).

The reduction of the risky win-stay bias did not reflect changes in the baseline probability of choosing safe (Figure 5 - figure supplement 1), and was observed on a rat-by-rat basis (pooling CaMKIIα-eNpHR3.0 and Pvalb-iCre-ChR2 rats, p=0.015; paired t-test across rats). Therefore, photoinhibition of lOFC specifically eliminated risky biases and not spatial biases. Moreover, risky biases were not sensitive to ITI duration (Figure 5C, sham), whereas spatial biases decreased with ITI duration (Figure 5B, sham), further suggesting that these sequential dependencies are dissociable.

## Discussion

Across species, OFC is critical for adapting to changing reward contingencies (Dias et al., 1996; Fellows and Farah, 2003; Schoenbaum et al., 2002), and value representations in OFC may drive learning (Miller et al., 2018). Our data are consistent with a role for OFC in updating rats’ risk attitudes or their beliefs about the world (Jones et al., 2012; McDannald et al., 2011; Miller et al., 2017; Wilson et al., 2014). lOFC neurons reflected reward history most strongly at trial initiation, when the animal has no information yet about the prospects on the current trial (Nogueira et al., 2017). These dynamics are therefore distinct from relative value coding observed in primate OFC, in which neurons encode the value of rewards on the current trial relative to rewards on the previous trial (Kennerley et al., 2011; Padoa-Schioppa, 2009; Saez et al., 2017). The temporal response profiles we observed are generally consistent with reports that OFC neurons fire transiently when an animal initiates reward-seeking behavior, here, trial initiation (Moorman and Aston-Jones, 2014).

We observed many “side-selective” neurons whose activity reflected which side the rat chose. A hallmark of primate OFC is that neurons do not encode spatial location (Grattan and Glimcher, 2014; Padoa-Schioppa and Cai, 2011). This discrepancy could reflect a species difference (Feierstein et al., 2006; Roesch et al., 2006). Alternatively, side-selectivity could reflect encoding of the left and right prospects or “goods” on each trial (Padoa-Schioppa, 2011).

We found that perturbation effects were uncoupled from task variables that seemed to be encoded most strongly, quantified by either fraction of neurons or discriminability, both during the cue period (reward history) and during the choice reporting period (left/right choice). It is possible that representations of reward history and choice may be distributed broadly enough that other brain areas can compensate for local perturbations, whereas representations used to update risk preferences may be more narrowly localized within or read out from OFC.

Are specific subcircuits within OFC responsible for the risky win-stay bias, and updating dynamic risk preferences more generally? The relatively small fraction of neurons (∼9%) that reflect whether choices are risky or safe may be behaviorally relevant, perhaps occupying a privileged position in the circuit and/or projecting to a common target. Alternatively, the more substantial fraction of neurons representing whether animals received reward (∼31%) may be involved in updating risk preferences, in which case their activity is read out to specifically update task-specific (here, risky), but not spatial, biases. This latter hypothesis is consistent with a recent study of medial OFC (mOFC) in mice (Namboodiri et al., 2019). In a Pavlovian conditioning paradigm, in which a tone probabilistically predicted a sucrose reward, mice exhibited trial-by-trial updating of their reward expectation, revealed by their anticipatory licking, based on reward history. Optogenetic inactivation of mOFC neurons projecting to the ventral tegmental area (VTA) during the reward period, but not the cue period, disrupted this trial-by-trial learning (Namboodiri et al., 2019). While our study targeted the lOFC in rats, the functional differences between medial and lateral OFC in rodents is not well understood, and our data are consistent with these results from mouse mOFC.

The risky win-stay bias may reflect evolutionary pressures in dynamic foraging environments, in which sequential successful outcomes are often not independent but reflect “clumped” resources (Blanchard et al., 2014; Wilke and Clark Barrett, 2009). However, while the risky win-stay bias may be an evolutionary relic of adaptive foraging behavior, today it demonstrably (and often adversely) affects behavior in finance, recreational gambling, and sports (Croson and Sundali, 2005; Hoffmann et al., 2010; Neiman and Loewenstein, 2011b). Our data show that this particular sequential bias is observable and manipulable in populations of neurons in lOFC.

OFC has been proposed to represent the animal’s location in a cognitive map of the task, which, in a reinforcement learning framework, corresponds to the current state in an abstract representation of task states and transitions between them (Nogueira et al., 2017; Wilson et al., 2014). The cognitive map hypothesis parsimoniously accounts for the results of OFC lesions in a variety of paradigms including delayed alternation, extinction, devaluation, and reversal learning, and is consistent with OFC’s role in evaluative processes such as regret (Gallagher et al., 1999; Izquierdo et al., 2004; Pickens et al., 2003; Steiner and Redish, 2014, 2012; Wilson et al., 2014). A recent study showed that the OFC may be particularly important for *learning* of action-outcome values (Miller et al., 2018). Our data are consistent with this hypothesis, and indicate that the coordinate space of the cognitive map in which OFC promoted learning was in task-specific (risky or safe), but not spatial (left or right) coordinates.

## Acknowledgements

We thank Paul Glimcher, Kenway Louie, Mike Long, David Schneider, Ben Scott, Emily Dennis, Mikio Aoi, Matthew Lovett-Barron, Cristina Domnisoru, Alejandro Ramirez and members of the Brody lab for helpful discussions and comments on the manuscript. We thank Alex Williams for feedback on the manuscript and for providing guidance and software for implementing TCA/PARAFAC tensor decomposition analysis. We thank Claudia Farb, Adam Carter, and Mike Hawken for reagents and assistance with histology and confocal imaging. We thank J. Teran, K. Osorio, L. Teachen, and A. Sirko for animal training. This work was funded in part by a K99/R00 award from NIMH (MH111926, to C.M.C.).

## Author Contributions

All authors provided feedback on analyses and the manuscript. C.M. Constantinople designed and performed all experiments, analyzed the data, and wrote the paper. A.T. Piet provided guidance for modeling and analysis. P. Bibawi performed histology. C.D. Kopec contributed to optogenetic experiments. A. Akrami and C.D. Kopec performed electrophysiological recordings to confirm photoinhibition from optogenetic constructs.

## Declaration of Interests

The authors declare no competing interests.

