## Supplemental Information for "Orbitofrontal cortex promotes trial-by-trial learning of risky, but not spatial, biases"

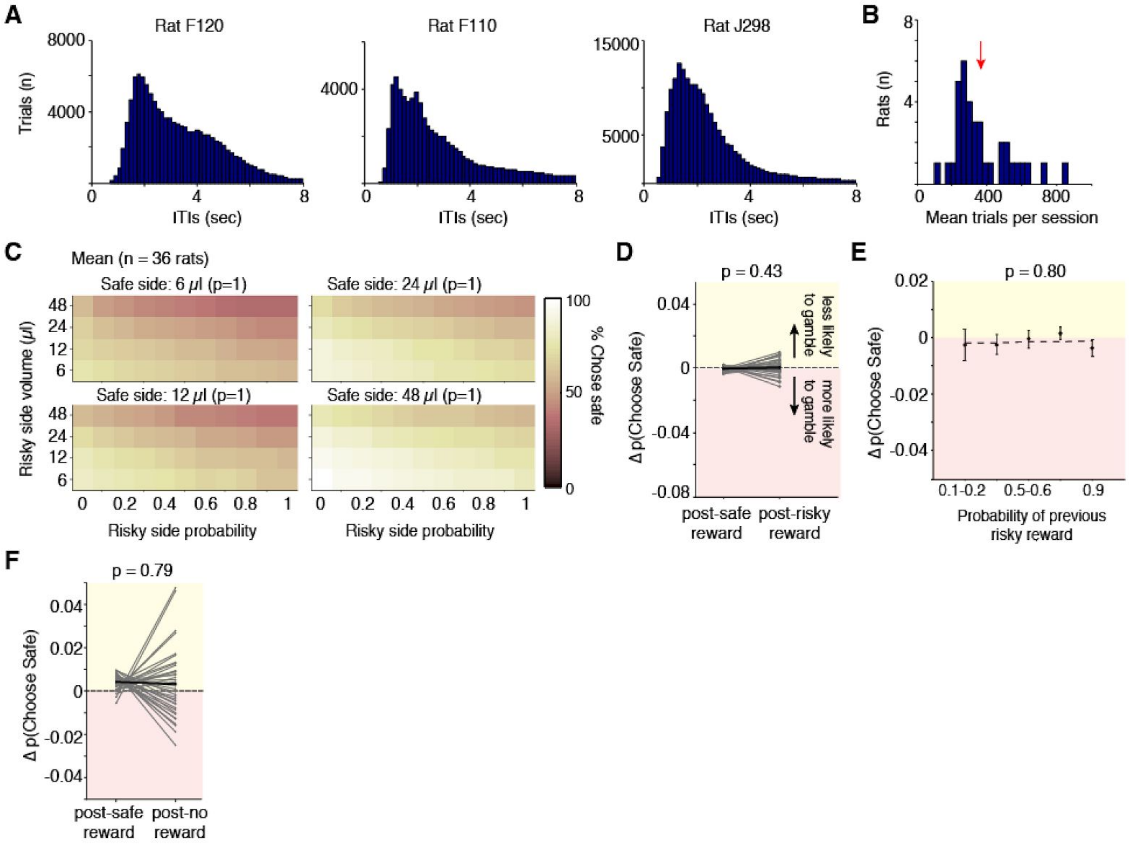


**Figure 1 - figure supplement 1.**

(A) Distribution of inter-trial intervals (ITIs) for three representative rats. Trials were self-paced, rats were free to initiate trials within 100-200ms of the preceding trial. If rats terminated the trial early by breaking center fixation, they were penalized with a time-out penalty (those trials are not shown).

(B) Average number of trials per session for each rat, excluding trials that were terminated prematurely. Mean of this distribution (368 trials/session) is shown by the red arrow.

(C) Mean behavioral performance across rats, including >2.5 million trials. Percent of trials all rats chose the safe option for each of the four safe side volumes. Axes show the probability and volume of risky alternatives. Mean performance across 36 rats (normalized to max before averaging).

(D) Estimates of conditional probabilities in finite sequential data can have small biases (Miller and Sanjurjo, 2015). If this bias were driving sequential effects in our data, such as increased willingness to take risks following risky wins, we reasoned that computing this bias from random flips (of the same length as our data) of a weighted coin would also reveal an effect. Therefore, we generated random choices for each rat with a generative probability corresponding to the mean probability of choosing the safe option for that rat. We then calculated the change in the probability of choosing the safe option based on reward history for the simulated choices; the same number of trials that were used in Figure 2C were applied to this analysis. There was no observable risky win-stay bias in the simulated dataset, indicating that the effect we observed did not reflect biased estimates of conditional probabilities.

(E) Difference in probability of choosing the safe option following guaranteed rewards and risky rewards of different probabilities (relative to the mean probability of choosing safe) for simulated data, as in B**.** Randomly simulated choices with the same sample sizes as the data (Figure 2D) did not exhibit a bias for risky choices with a graded dependence on reward probability. $p = 0.80$ of slope parameter of least-squares regression line (dashed line). Therefore, the risky win-stay bias we observe, with graded dependence on reward probability, does not reflect biased estimation of conditional probabilities.

(F) Difference in probability of choosing the safe option following guaranteed rewards, or risky unrewarded choices. There was no systematic, significant change in probability of choosing safe following unrewarded trials (paired t-test comparing change in probability of choosing safe).

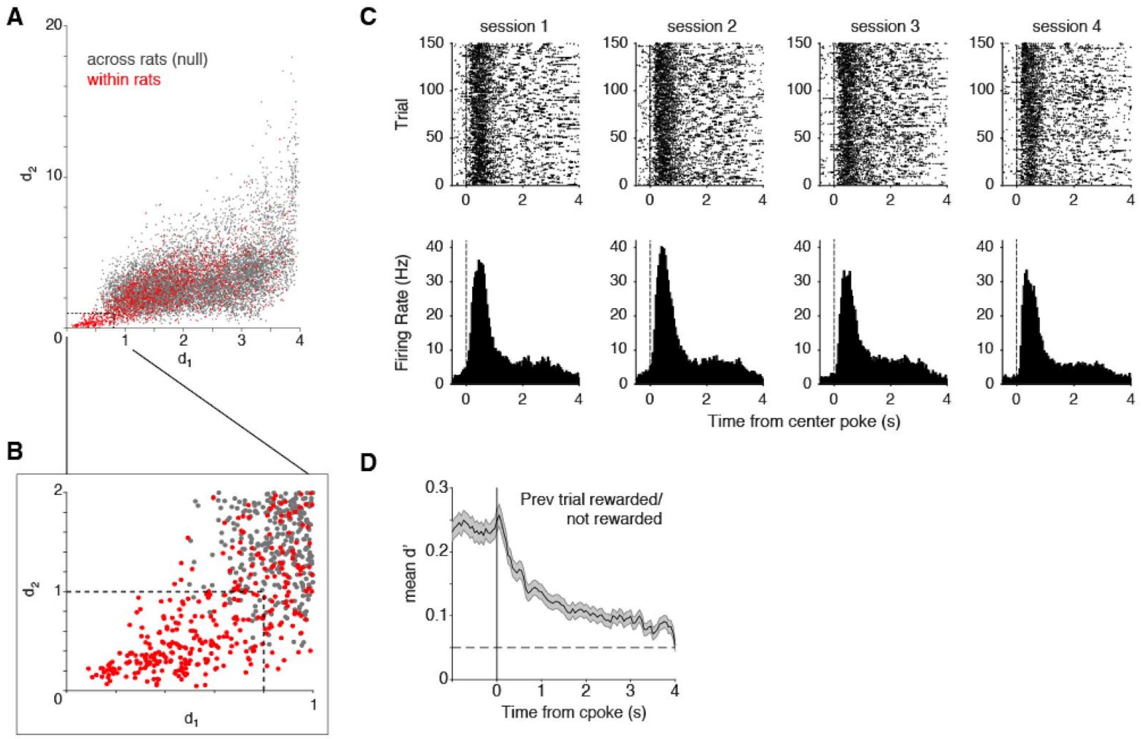


**Figure 2 - figure supplement 1.**

(A) Distribution of $d_1$ and $d_2$ values comparing waveforms across rats produces a null distribution (gray, see Methods). Distribution of values comparing waveforms within rats from subsequent recording sessions (red). Dashed lines are empirically chosen thresholds.

(B) Subplot from panel A.

(C) Neuron that was identified as putatively identical across four recording sessions. Raster plots only show 150 trials (out of 400-600 each day) for display purposes (upper panels). PSTHs (derived from all trials) are shown below (lower panels).

(D) Figure 3B was reproduced combining putatively identical units recorded over multiple days. Mean discriminability index (d') depending on whether the previous trial was rewarded or not, computed in 50ms bins. Error bars are +/- s.e.m.

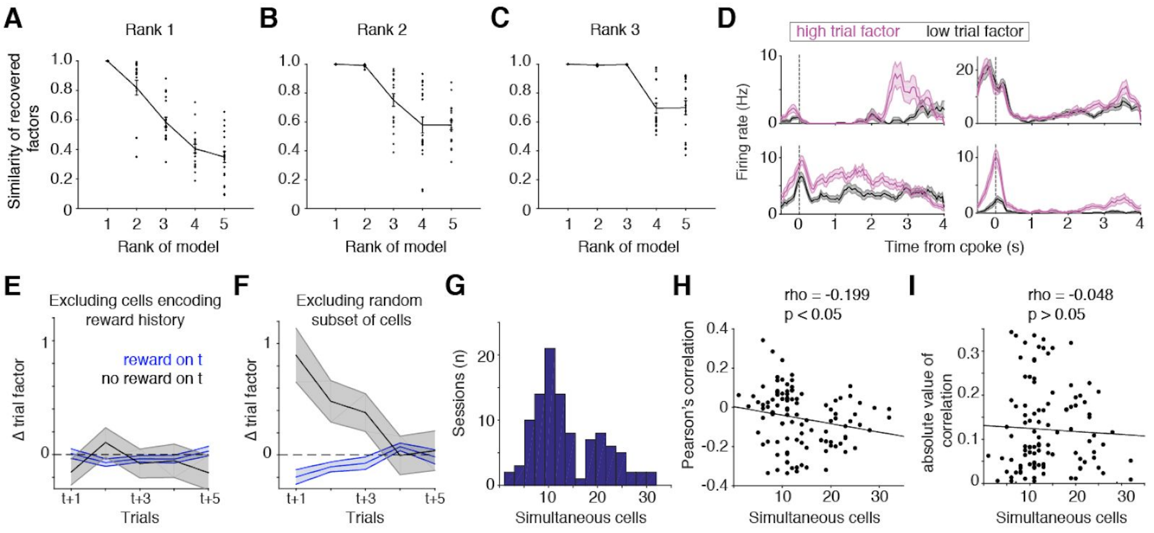


**Figure 2 - figure supplement 2.**

(A-C) Method used to determine model rank. We performed 20 random initializations and compared the similarity of the factors recovered from each iteration to those recovered from the previous one. We show the distribution of similarity indices for example recording sessions that were determined to be rank 1, 2, and 3 (A,B,C, respectively). Black lines are mean +/- s.e.m. The majority of the data was either rank 1 or 2 (50/105 sessions were rank 1, 50/105 were rank 2, 5/105 were rank 3), so for simplicity, we fit a rank 1 model to each session.

(D) Four neurons from the recording session in panel 3D; firing rates are plotted when the trial factor was high (>85th percentile) or low (<15th percentile).

(E) Mean (+/- s.d.) shuffle-corrected reward (blue) and no-reward (black) triggered averages of trial factors across all sessions (see Methods), excluding cells that had significantly different spike counts following rewarded or unrewarded trials.

(F) Mean (+/- s.d.) shuffle-corrected reward (blue) and no-reward (black) triggered averages of trial factors, excluding random subsets of cells, of the same number that were excluded in panel E.

(G) Distribution of simultaneously recorded units across all recording sessions.

(H) Relationship between the Pearson's correlation between trial factors and reward history, and number of units recorded in each session.

(I) Relationship between the absolute magnitude of the Pearson's correlation between trial factors and reward history, and number of units recorded in each session.

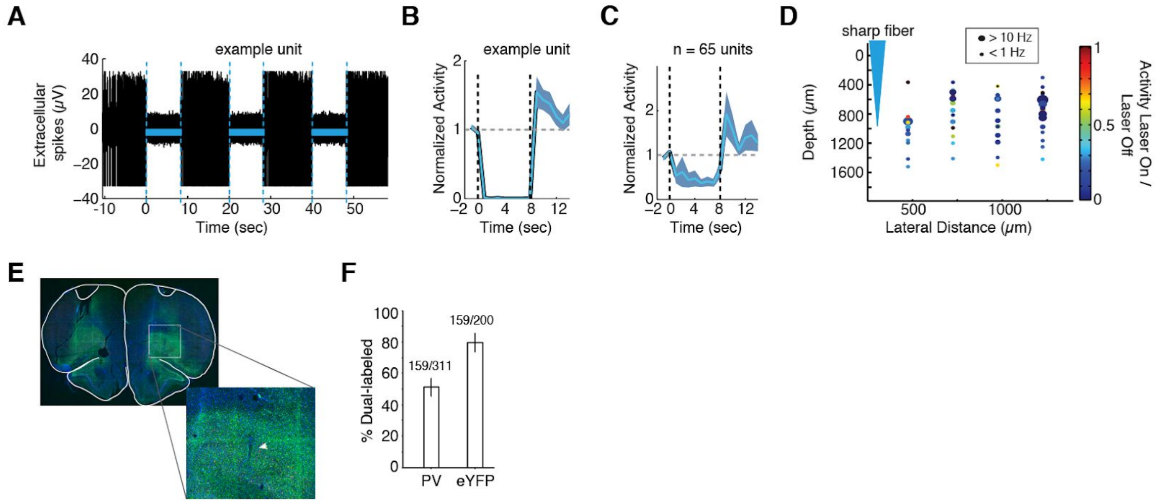


**Figure 3 - figure supplement 1.**

(A) Representative unit recorded from Pvalb-iCre rats expressing ChR2. Three epochs of photoinhibition (blue lines) reliably suppressed spiking activity. Blue laser was pulsed at 20Hz, 10ms pulse width, for 8 seconds.

(B) Normalized activity of the cell shown in A; mean +/- s.e.m. over 30 photoinhibition epochs.

(C) Mean suppression over 65 recorded units from 2 rats.

(D) Activity change of each unit plotted as a function of its distance from the optical fiber. Units were recorded in four tracks, 250, 500, 750, or 1000 μm from the fiber tip. Robust photoinhibition was observed in all tracks.

(E) Example injection site shown in Figure 4C; inset shows putative fiber track.

(F) Percent of parvalbumin-immunoreactive cells that co-expressed eYFP in Pvalb-iCre rats expressing eYFP-ChR2 (left), and fraction of eYFP-expressing cells co-labeled for parvalbumin immunoreactivity (right).

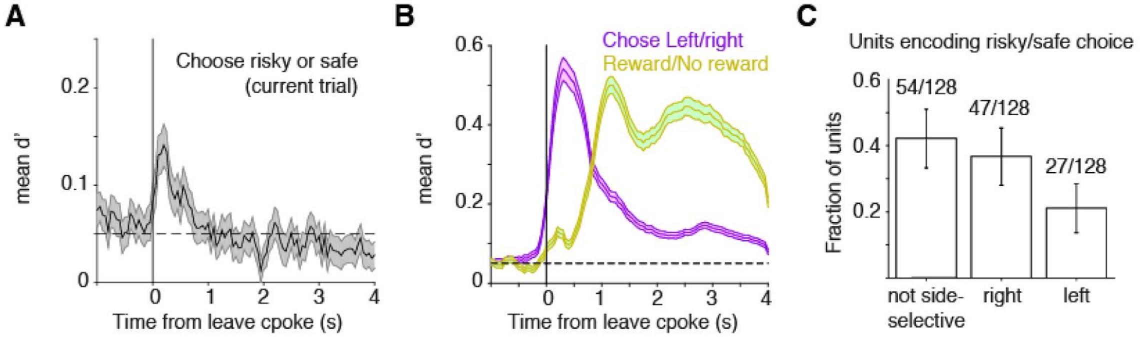


**Figure 4 - figure supplement 1.**

(A,B) Figure 4B (A) and 4F (B) were reproduced combining putatively identical units recorded over multiple days. Mean discriminability index (d') depending on whether the rat chose the safe or risky option on rewarded trials only (A), chose left or right (B, purple), or was rewarded (B, yellow), computed in 50ms bins. Error bars are +/- s.e.m.

(C) Of the units with significantly different spike counts on trials in which rats chose risky or safe, the fraction selective (or not) for choosing the left or right port.

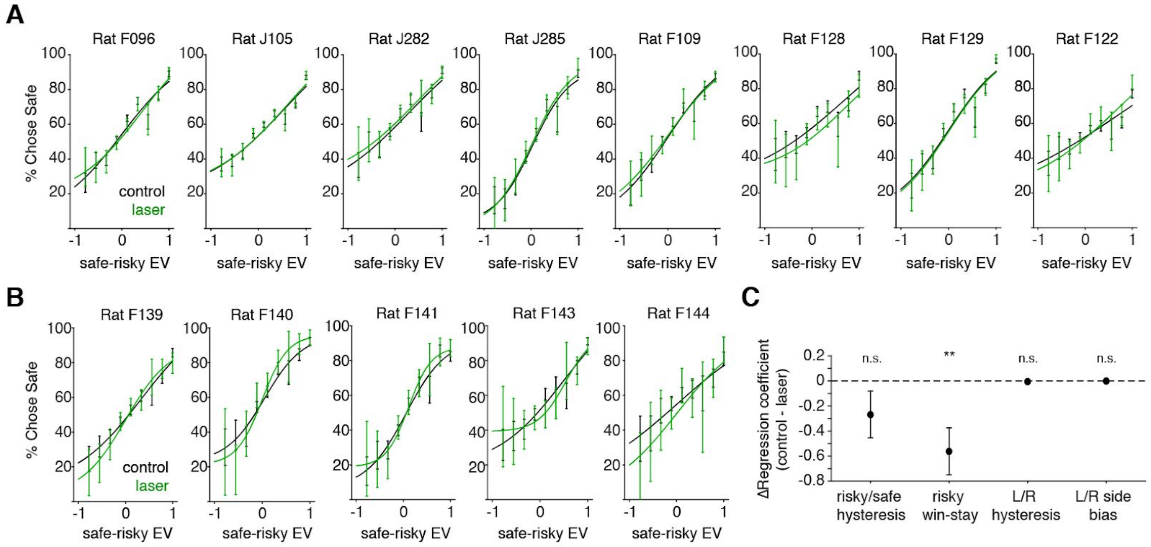


**Figure 5 - supplement 1.**

(A) Psychometric performance for each CaMKIIα-eNpHR3.0 rat on control trials (black) and trials following photoinhibition during the choice period. Rat J285 is replotted from Figure 6. These plots include all trials, regardless of trial history, so the elimination of the risky win-stay bias is not evident.

(B) Psychometric performance for each Pvalb-iCre-ChR2 rat on control trials (black) and trials following photoinhibition during the choice period.

(C) Difference in logistic regression coefficients (control - photoinhibition) parameterizing different choice biases. Data are mean +/- standard deviation across rats. Asterisks indicate significant Bonferroni-corrected p-value from one-way ANOVA (p=0.0063).

### Methods

#### Subjects

A total of 39 male rats between the ages of 6 and 24 months were used for this study. These included 35 Long-evans and 4 Sprague-Dawley rats (*Rattus norvegicus*). Of these, 3 rats were used for neural recordings, and 16 for optogenetic experiments, including LE-Tg (Pvalb-iCre)2Ottc rats (n=5) made at NIDA/NIMH and obtained from the University of Missouri RRRC (transgenic line 0773). These are BAC transgenic rats expressing Cre recombinase in parvalbumin expressing neurons. Investigators were not blinded to experimental groups during data collection or analysis. Animal use procedures were approved by the Princeton University Institutional Animal Care and Use Committee and carried out in accordance with National Institutes of Health standards.

Animals were water restricted to motivate them to perform behavioral trials. They obtained water rewards during behavioral training sessions, which ranged from 1-5 hours per day, and an ad lib access period of up to 1 hour. Food was typically placed in the behavioral box during training, so it was available during the water access period. All rats obtained a minimum volume of water equal to 3-5% of their body mass, (30-50 mL/kg). Water consumption was monitored during the behavioral session, and if rats consumed less than the minimum requirement, additional water was offered during an ad lib period. The ad lib period terminated either when the target water volume was exceeded or after 1 hour.

#### Behavior

We have previously described rats' behavior on this task in detail (Constantinople et al., 2019). Briefly, rats were trained in a high-throughput facility using a computerized training protocol. Rats were trained in operant training boxes with three nose ports. When an LED from the center port was illuminated, the animal could initiate a trial by poking his nose in that port; upon trial initiation the center LED turned off. While in the center port, rats were continuously presented with a train of randomly timed clicks from a left speaker and, simultaneously, a different train of clicks from a right speaker. The click trains were generated by Poisson processes with different underlying rates (Hanks et al., 2015); the rates conveyed the water volume baited at each side port. After a variable pre-flash interval ranging from 0 to 350ms, rats were also presented with light flashes from the left and right side ports; the number of flashes conveyed reward probability at each port. Each flash was 20ms in duration; flashes were presented in fixed bins, spaced every 250ms, to avoid perceptual fusion of consecutive flashes. After a variable post-flash delay period from 0 to 500ms, the end of the trial was cued by a go sound and the center LED turning back on. The animal was then free to choose the left or right center port, and potentially collect reward.

In this task, the rats were required to reveal their preference between safe and risky rewards. First, rats proceeded through a series of early training stages included training the rat to center poke, gradually growing the duration of center fixation, and introducing cues representing certain rewards of each volume on one side at a time. Once they were in the final training stage they were presented with the full stimulus set. To determine when rats were sufficiently trained to understand the meaning of the cues in the task, we evaluated the "efficiency" of their choices as follows. For each training session, we computed the average expected value per trial of an agent that chose randomly, and an expected value maximizer, or an agent that always chose the side with the greater expected value. We compared the expected value per trial from the rat's choices relative to these lower and upper bounds. Specifically, the efficiency was calculated as follows:

$$efficiency = 0.5\frac{rat_{EV/trial} - rand_{EV/trial}}{EVmax_{EV/trial} - rand_{EV/trial}} + 0.5$$

The threshold for analysis was the median performance of all sessions minus 1.5 times the interquartile range of performance across the second half of all sessions. Once performance surpassed this threshold, it was typically stable across months. Occasional days with poor performance were usually due to hardware malfunctions in the rig or a change in the experiment (e.g., the first day being tethered for electrophysiological recordings). Days in which performance was below threshold were excluded from analysis.

**Psychometric curves**

We measured rats' psychometric performance when choosing between the safe and risky options. For these analyses, we excluded trials where both the left and right side ports offered certain rewards. We binned the data into 11 bins of the difference in the expected value (reward x probability) of the safe minus the risky option. Psychometric plots show the probability that the subjects chose the safe option as a function of this difference (see Figure S1C). We fit a 4-parameter sigmoid of the form:

$$p(Choose_S) = y_0 + \frac{1-2a}{(1+e^{(-b(V_S - V_R - x_0))})},$$

where $y_0$, $a$, $b$, and $x_0$ were free parameters. Parameters were fit using a gradient-descent algorithm to minimize the mean square error between the data and the sigmoid, using the sqp algorithm in Matlab's constrained optimization function fmincon.

**Chronic electrophysiology**

Tetrodes were constructed from twisted wires that were either PtIr (18 µm, California Fine Wire) or NiCr (25 µm, Sandvik). Tetrode tips were platinum- or gold-plated to reduce impedances to 100-250 kΩ at 1kHz using a nanoZ (White Matter LLC).

Microdrive assemblies were custom-made as described previously (Aronov and Tank, 2014). Each drive contained 8 independently movable tetrodes, plus an immobile PtIR reference electrode. Each animal was implanted over the right OFC. On the day of implantation, electrodes were lowered to ~4.1 mm DV. Animals were allowed to recover for 2-3 weeks before recording. Shuttles were lowered ~30-60 µm approximately every 2-4 days.

Data was acquired using a Neuralynx data acquisition system. Spikes were manually sorted using MClust software. Units with fewer than 1% inter-spike intervals less than 2ms were deemed single units. All units that fired more than two spikes on half of trials were included in analysis (n=1459/1881). To convert spikes to firing rates, spike counts were binned in 50 ms bins and smoothed using Matlab's smooth.m function.

**Discriminability, or *d'*, of OFC neurons**

To measure neuronal discriminability for different task variables, such as whether the previous trial was rewarded, we computed the mean difference in the smoothed firing rate on different trial types divided by the square root of their mean variance:

$$d' = \frac{|\mu_1 - \mu_2|}{\sqrt{\frac{1}{2}(\sigma_1^2 + \sigma_2^2)}}.$$

Because we computed the absolute value of the difference in firing rates, we subtracted the mean shuffled *d'*, computed from shuffling the data 15 times. *d'* was computed in 50 ms bins, as this was the bin-width used for computing the firing rates (see above).

**Spike waveform analysis for identifying the same neurons recorded over days**

The single neuron data shown in Figures 3 and 4 treated each unit recorded on a different day/recording session as a unique unit. However, we also modified previously published methods (Tolias et al., 2007) to identify units recorded over days (Figure S2). We computed two metrics (Tolias et al., 2007), which we describe below, based on the spike waveform. The first metric compared how similar the shape of the waveform was across recording sessions. For each waveform on session 1 ($x$), we computed $\alpha$ to make it as close as possible to the waveform on session 2 ($y$):

$$\alpha(x, y) = \underset{\alpha}{\operatorname{argmin}} ||\alpha x - y||^2.$$

We used Matlab's constrained minimization function fmincon.m to find $\alpha$. We then computed the Euclidean distance between the scaled waveforms, $d_1$.

$$d_1(X,Y) = \sum_{i=4}^{4} \frac{||\alpha(x_i, y_i)x_i - y_i||}{||y_i||}$$

The second metric, $d_2$ quantified the difference in amplitude across the 4 channels of each tetrode.

$$d_2(X,Y) = \max_{i=4}^{4} |log(\alpha(x_i, y_i))| + \max_{i,j}^{4} |log(\alpha(x_i, y_i)) - log(\alpha(x_j, y_j))|$$

We computed these metrics for all pairs of waveforms recorded on the same tetrode on subsequent recording sessions. To compare these values to a null distribution, we computed the $d_1$ and $d_2$ metrics for units recorded from two different animals, which could not have identical waveforms. We used this null distribution to empirically determine thresholds for $d_1$ (0.8) and $d_2$ (1). Units recorded on consecutive sessions with values below these thresholds, and with significant Pearson's correlation coefficients of their mean firing rates aligned to trial start ($p<0.05$), were tentatively classified as identical (Figure S2A-C). Putatively identical neurons were then manually examined, and those that exhibited qualitatively different PSTHs, or different mean firing rates over days were rejected and treated as separate units. 191/1459 (13%) units that met criteria for inclusion in analysis were recorded over multiple sessions. Combining data from these units over sessions did not change the results (Figure S2D-G). For population analyses (TCA/PARAFAC; Figure 3E-G), data were not combined across recording sessions.

**Tensor components analysis/CANDECOMP/PARAFAC tensor decomposition**

To fit the tensor decomposition model, we used software recently made publicly available (Williams et al., 2018): https://github.com/ahwillia/tensortools.

To initially determine the dimensionality, or rank, that should be applied to each recording session, we iteratively tried different numbers of dimensions, or "tensor components", and computed a similarity index to determine how sensitive the recovered factors were to the initialized values of the optimization procedure (Williams et al., 2018). The similarity index was computed on factors recovered from consecutive initializations using the score.m function in Matlab's Tensor Toolbox. The maximum number of components that yielded an average similarity index >90% was used as the number of components, or rank, for each recording session (Figure S3). Nearly all of our recording sessions were rank 1 or 2 (by this method). TCA/PARAFAC is notably different from principal components analysis (PCA) in that the first component does not necessarily explain the most variance of the data (Williams et al., 2018). Therefore, given that most of our data was low rank, to simplify the problem of determining which trial factors to analyze, we fit a rank 1 model to each recording session.

We computed the shuffle-corrected reward-triggered averages of trial factors (and no reward-triggered averages) as follows. We computed the average change in trial factors relative to the mean trial factor relative to each rewarded (and unrewarded) trial, up to 7 trials in the future. We then performed a shift correction, shuffling the trials randomly with respect to reward history, and computed the average change in trial factors relative to rewarded and unrewarded trials (relative to the mean) for the shuffled data. We subtracted the shuffled averages from the true averages to obtain the plot in Figure 3F.

TCA decomposes a 3rd order data tensor $X_{n,t,k}$ (with n neurons over k trials of length t) by a sum of rank 1 factors $\sum_{r=1}^{R} w_r b_r a_r$. Here, for each rank r, w is a vector of neuron factors, b is a vector of temporal (time within trial) factors, and a is a vector of across trial factors. We note that each of the factors (neuron, temporal, and trial) is a linear gain factor, multiplied by the others; therefore, meaningful units are difficult to determine (as all the factors are multiplied); their scale is a gain on the other terms.

**Acute electrophysiology**

To confirm photoinhibition in Pvalb-iCre-ChR2 rats, we performed virus delivery as described above. After 6-8 weeks to allow for expression, rats (n=2) were anesthetized with for surgery with 0.2mL ketamine and 0.2mL buprenorphine. Craniotomies were made over frontal orienting field (FOF; centered 2 mm anterior to the Bregma and 1.3 mm lateral to the midline), and the rat was maintained under isoflurane anesthesia.

A chemically sharpened fiber optic (50um core, 125um cladding) was inserted into the field of infected neurons to a depth of 1mm. A sharp tungsten electrode (0.5MW) mounted to a Narishige oil hydraulic micromanipulator was manually lowered into the brain. Recordings were made using a Neuralynx Cheetah system applying a bandpass filter from 300-6000 Hz to the voltage signal. At each site an 8s laser illumination (473nm, 25mW, 20Hz, 20ms of pulse duration) was delivered every 20s, 10 times. A mechanical shutter (Thor Labs optical beam shutter) was used to control the laser timing.

Spikes were automatically detected as brief (<1ms) events that crossed a threshold of +/- 5 standard deviations on the filtered voltage trace. In order to remove light artifact and possible population spikes of driven PV neurons, we removed spikes within the 20ms of the laser pulse onset. Inhibition was defined as the mean spike rate during the 8s laser on period / the mean spike rate during the 12s laser off period.

**Optical fiber chemical sharpening**

We used standard off the shelf FC-FC duplex fiber optic cables (#FCC2433, FiberCables.com), and as previously described(Hanks et al., 2015), stripped the outer plastic coating. To etch the fiber, 2-2.5 mm of the fiber tip was submerged in 48% hydrofluoric acid with mineral oil on top. Over the course of ~17 minutes, a motor (Narishige) slowly pulled the fiber tip out of the hydrofluoric acid, producing a long taper. The speed of the motor was then increased and maintained at a constant speed until the tip was entirely removed from the acid (usually by 13-15 minutes). This protocol reliably produced sharp, well-etched fibers with uniform and broad light scatter. Fibers that did not produce sufficiently broad or uniform scatter were discarded.

**Virus delivery and fiber implantation**

We used methods described previously (Hanks et al., 2015); here we describe procedures specific to this experiment. We injected 2 µL of AAV virus (AAV5-CaMKIIα-eNpHR3.0-eYFP in wild type rats, or AAV-FLEX-rev-ChR2-tdtomato in Pvalb-iCre rats) using a Nanoject (Drummond Scientific). Six closely spaced injection tracts (typically 500 µm apart) were made in each craniotomy; each rat had bilateral craniotomies and injections, so there were 12 total injection tracts per animal. For OFC injections, in each track, 18 injections of 14.1 nL were made every 100 µm in depth starting at 3.7 mm below brain surface (3.7-5.4 mm DV). Virus was expelled at 20nL/sec. Injections were made once every 10 seconds; at the final injection in a tract, the pipette was left in place for at least 2 minutes before removal.

Chemically sharpened fibers (50 µm core, 125µm cladding) were implanted at 5° angles relative to the midline. Fiber tips were positioned 0.4 mm lateral from the center track at brain surface so that, when the tip was lowered to 4.6 mm DV, it was centered at the target coordinates (+3.5 AP, +/- 2.5 ML). Viral constructs were allowed to develop for 6-8 weeks before behavioral experiments began.

**Optogenetic perturbation**

For bilateral halorhodopsin inactivations, the laser beam from a 200 mW, 532 nm laser (OEM Laser Systems) was split into two beams of roughly equal power (~25 mW) using a beam splitter (Doric DMC_1x2i_VIS_FC). Laser illumination was delivered on a subset of trials by opening a shutter with a 5 V TTL. On a random subset of 15% of trials, illumination occurred during the entire trial (triggered when the rat entered the center poke until he was free to leave it), or on a random 15% of trials, illumination was triggered when the rat left the center poke, and persisted for the first 4 seconds of the inter-trial interval, resulting in transient silencing of cortical dynamics. In Pvalb-iCre-ChR2 rats, we used a 473 nm laser. Laser pulses (10ms pulse width) were delivered at 20 Hz.

The rats generally chose risky on a minority of trials, and by definition, the fraction of those that were rewarded is a smaller subset. Moreover, rewarded trials were more frequent than non-rewarded trials. To overcome these challenges, we collected data from many sessions. The median number of sessions in which rats experienced photoinhibition was 27 (range: 21-70, mean: 37). We did not observe differences in the optogenetic result when comparing the first or second half of all sessions (data not shown).

**Normal approximation of 95% binomial confidence intervals**

To compute error bars on the choice probabilities for the risky and spatial win-stay biases (Figures 3J, 4J, 4L), we computed the 95% confidence intervals using the test statistic for the chi-square distribution (Corder and Foreman, 2014) as follows:

$$CI_{95} = z\sqrt{\frac{p_{psr}(1-p_{psr})}{n_{psr}} + \frac{p_{prr}(1-p_{prr})}{n_{prr}}}$$

where $p_{psr}$ is the probability of choosing safe following a safe reward ("post-safe reward") and $p_{prr}$ is the probability of choosing safe "post-risky reward", $n$ is the number of observations for each condition (post-safe reward and post-risky reward), and $z$ the z-score for 95% confidence intervals from a normal distribution.

**Logistic regression model of choice biases**

To evaluate the contribution of different potential choice biases to behavior, we implemented a logistic regression model, in which we parameterized the rats' probability of choosing right as follows:

$$p(ChooseR) = 1/(1+exp(\Delta EV + rs_{hyst} + risky_{win-stay} + lr_{hyst} + lr_{bias}))$$,

where $\Delta EV$ is the right minus left expected value (reward x probability) on each trial, $rs_{hyst}$ captures risky/safe hysteresis (i.e., if the rat just chose safe, the likelihood he will choose safe on the next trial), $risky_{win-stay}$ parameterizes increased willingness to choose the gamble conditioned on a risky win, $lr_{hyst}$ parameterizes the probability of repeating left/right choices, and $lr_{bias}$ parameterizes overall side biases for the left and right port. The only parameter that was significantly changed (and in fact reduced) by photoinhibition during the choice report was the $risky_{win-stay}$ parameter (p=0.0063, one-way ANOVA comparing parameters fit to control and opto conditions across rats, Bonferroni correction for multiple comparisons).
